# Wide-Field Multicontrast Nonlinear Microscopy for Histopathology

**DOI:** 10.1101/2022.04.16.488489

**Authors:** Leonardo Uribe Castaño, Kamdin Mirsanaye, Ahmad Golaraei, Lukas Kontenis, Susan J. Done, Vuk Stambolic, Margarete K. Akens, Brian C. Wilson, Virginijus Barzda

## Abstract

A multicontrast polarimetric wide-field second harmonic generation (SHG) and multiphoton excitation fluorescence (MPF) microscope is optimized for large area imaging of hematoxylin and eosin (H&E) stained and unstained histology slides. The bleaching kinetics of MPF and SHG are examined with various laser intensities at different pulse repetition rates to determine the optimal wide-filed imaging conditions for H&E stained histology slides. Several polarimetric parameters are used to investigate the organization of extracellular matrix collagen in the histology samples.

## 1. Introduction

Digital pathology has been adopted for cancer diagnostics and relies on a whole slide scanning of histology samples. [1–7]. Most often hematoxylin and eosin (H&E) stained histology slides are used as a gold standard for cancer diagnostics and are typically imaged with white-light or fluorescence image contrasts [8]. Multicontrast nonlinear optical microscopy with multiphoton excitation fluorescence (MPF), second harmonic generation (SHG) and third harmonic generation (THG) can also be employed for histopathology investigations [9–19]. Imaging the same slides with nonlinear microscopy presents a convenient way of gathering more information about the tissue structure without altering the H&E slide preparation procedure. Therefore, the new and retrospective histopathology investigations can be readily implemented with nonlinear microscopy. Multiphoton excitation fluorescence in H&E stained tissue slides is provided predominantly by eosin dye fluorescence, which highlights various cellular proteins and extracellular matrix [9]. The second harmonic generation image contrast is particularly suited for visualizing collagen [20, 21]. SHG is sensitive to polarization, therefore ample of information can be extracted from polarimetric measurements about collagen ultrastructure, which often is affected by tumorigenesis [13, 14, 19, 22–32]. Hence, multicontrast imaging of H&E slides provides complementary information that can be used for histopathology studies.

Whole slide imaging is needed for digital pathology investigations. Image of a whole slide is acquired by tiling smaller images, or by a wide line scanning. Nonlinear optical microscopy is typically performed using a laser scanning microscope that allows for optical sectioning; however, scanning is rather slow when large area images have to be acquired. Nonlinear microscopy can also be performed in a wide-field imaging mode, which provides video rate imaging for a large sample area due to parallel detection with a camera compared to a sequential point by point detection in the scanning mode [33]. A limiting factor of the wide-field nonlinear microscopy is the reduced optical sectioning, however, the histopathology investigations are conducted on thin tissue sections (∼ 5 *μm*), therefore, the wide-field nonlinear microscopy is well-suited for the digital pathology.

For wide-field nonlinear microscopy, a high-power ultrafast laser is employed to provide enough pulse energy for a large area illumination. The illumination is achieved either by a high-power oscillator [34], an amplified laser system [35], or a fiber laser [36]. Wide-field illumination using high-powered lasers differently affects biological samples when compared to a raster-scanning with tightly focused laser beam. In the laser scanning mode, a laser beam is focused onto a diffraction limited focal volume and a sample area is scanned with pixel dwell time of tenths of microseconds. In the wide-field nonlinear microscopy mode, a laser beam illuminates whole field of view and the signal integration time depends on camera frame rate. The illumination time can last from several milliseconds to seconds. Simultaneous illumination of all pixels at once allows for much longer integration time per pixel, and therefore the intensity of illumination can be reduced. However, the high incident laser power may deposit substantial amount of energy into the sample, leading to adverse effects. These effects may induce sample photobleaching.

Photobleaching is particularly detrimental when prolonged illumination is required. This may happen when polarization measurements of SHG are performed for a longer period. Measurements with several incident, or incident and outgoing polarization states are required for ultrastructural characterization of tissue collagen [10, 20, 37]. Bleaching has to be well characterized when sequential measurements with different polarization states are performed and polarimetric parameters such as SHG circular dichroism (SHG_CD_), SHG linear dichroism (SHG_LD_) or polarization-in polarization-out (PIPO) measurements are employed [10, 38–41]. Therefore, for a wide-field nonlinear microscopy, a careful assessment of the laser illumination parameters and bleaching effects in the biological samples are required to optimize the imaging conditions and to avoid possible artefacts, in particular with the polarization measurements.

In this work, we examine the MPF and SHG signal stability for the wide-field nonlinear microscopy imaging of H&E stained and unstained histology samples. A theoretical and experimental description of fluorescence and SHG bleaching in H&E stained samples is provided, and the laser settings are determined for wide-field nonlinear optical imaging. An example of polarimetric wide-field SHG microscopy for a large area (700*μm* x 700*μm*) imaging of histology tissue microarray cores is presented. The newly developed wide-field polarimetric imaging method can be readily applied for nonlinear digital pathology investigations [42].

## 2. Materials and Methods

### 2.1. Wide-field polarimetric nonlinear optical microscope

The wide-field microscope is based on the previously described setup [33] with addition of polarization state generator and polarization state analyzer [42]. The setup outline is presented in Fig. 1. A high-power amplified KGW laser (Pharos, Light Conversion, Lithuania) is used for a large area wide-field illumination. The laser beam is collimated to ≈ 4mm in diameter and coupled to the microscope. The beam is passed through a polarizing cube and a liquid crystal variable retarder (Thorlabs, LCC1423-A), which constitutes the polarization state generator (PSG). An achromatic doublet with a focal length of 75 mm is used to focus the beam for sample illumination. In order to obtain wide-field imaging, the sample is placed above the focal plane. Therefore, the illumination area can be adjusted by axially translating the doublet. The sample with SHG signal generated in the forward direction is imaged by a 20× 0.5 numerical aperture (NA) air objective lens (Carl Zeiss, Germany). The polarization state analyzer (PSA) containing a liquid crystal retarder (Thorlabs, LCC1423-B) is inserted in the collimated path of the infinity corrected objective. The SHG signal then is passed through a tube lens, a linear polarizing cube, a dichroic mirror, and two filters (a BG39 Shott glass filter, and a 515 nm interference filter with 10 nm bandwidth). The filtered SHG signal is focused onto sCMOS camera (Hamamatsu Orca-Flash 4) containing sensor with 2056×2056 pixel array. For MPF imaging, the filters are exchanged to a 578nm bandpass filter (Semrock, FF01-578/105).

**Fig. 1.**
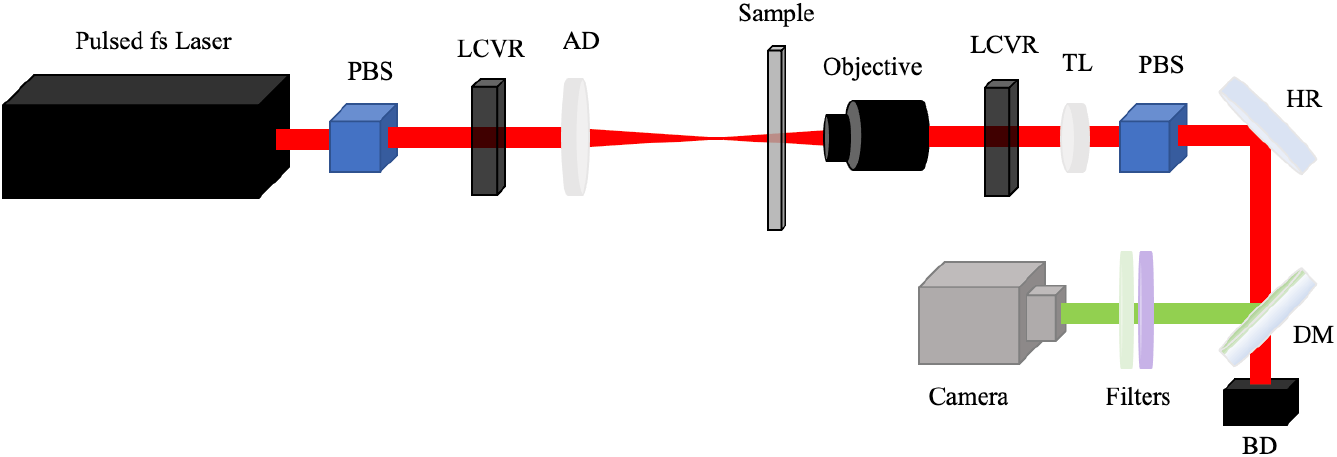
The outline of wide-field polarimetric nonlinear microscopy setup. The laser beam is passed through the polarization state generator containing polarizing beam splitter (PBS) and liquid crystal variable retarder (LCVR) that creates an appropriate polarization state for the fundamental beam. A 75mm focal length achromatic doublet (AD) focuses the beam just before the sample. The SHG or MPF and fundamental beams are collected by the Objective. The polarization state analyzer containing LCVR and a second PBS probe different polarization states of the SHG or MPF signal. The tube lens (TL) focuses the beam onto the camera. The beam passes high reflection mirror (HR), dichroic mirror (DM) and appropriate filters. The separated laser beam from the DM is directed to the beam dump (BD). The actual setup is built vertically in upright configuration.

The calibration of polarization setup was accomplished using Z-cut quartz plate and a thin lithium triborate (LBO) crystal of 0.1 mm thickness attached to a thin electron microscope gold grid. The SHG signal intensity had a Gaussian intensity profile along both lateral (horizontal - and vertical /) directions in the image.

### 2.2. Nonlinear excitation fluorescence bleaching kinetics

Bleaching of multiphoton excitation fluorescence is modeled by assuming photo-induced accumulation of non-fluorescing chromophores without further quenching of the fluorescing molecules by the photo-products. The fluorescence intensity ***I***_*f*_ is proportional to the two-photon absorption rate, σ, the fluorescence yield, Φ_*f*_ and the concentration of fluorophores,.*c*. The fluorescence is also proportional to the pulse energy per unit area squared, *U*^2^, for two-photon process, and the pulse repetition rate, *r*, [43]:

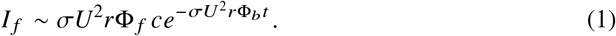

Details of the bleaching model are presented in the Appendix. Upon illumination, the fluorescence undergoes time, *t*, dependent exponential bleaching with rates proportional to the two-photon absorption rate, bleaching yield, Φ_*b*_, the pulse energy per area squared and the pulse repetition rate.

The measured fluorescence *I*_*f*_ normalized to the initial fluorescence *I*_0_ is fitted as a bi-exponential decay as follows:

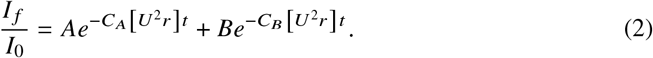

The bi-exponential decay indicates the presence of two or more fluorescing species present in the tissue, with heterogeneous population that could be approximated with the two-exponential bleaching kinetics. The amplitudes *A* and *B* represent the respective fraction of each fluorescing species, where *A* + *B* = 1. The *C*_*A*_ and *C*_*B*_ constants are the products of two-photon absorption cross-sections and bleaching yields for two chromophore types. The *C*_*A*_ and *C*_*B*_ constants can be extracted from fitting the bleaching decays. The weighted amplitude fluorescence bleaching rate Γ_*AB*_ can also be calculated as:

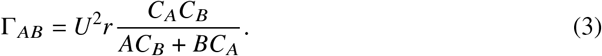

It can be seen that bleaching rate depends on the pulse energy per area squared and repetition rate of the pulses. The *C*_*A*_ and *C*_*B*_ are constants, and therefore should be similar for various values of pulse energy per area and pulse repetition rate. If the *C*_*A*_ and *C*_*B*_ appear very different for different excitation conditions, that indicates the presence of additional processes not accounted for by the bleaching model.

### 2.3. SHG Polarimetric analysis

The polarimetric measurements are performed by employing liquid crystal variable retarders (LCVR), one for PSG and one for PSA. Two perpendicular linear or circular polarization states can be produced with the LCVR oriented with fast axis at 45 degrees from the incident linear polarization orientation [15, 42]. The SHG signal intensity, *I*_*M*_, measured after the PSA containing a single LCVR oriented at 45 degrees with respect to vertical polarization can be characterized with:

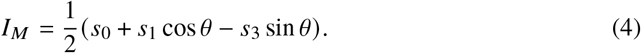

where *s*_0_, *s*_1_, *s*_2_, *s*_3_ are the measured SHG Stokes vector components [44, 45] and θ is the retardance angle of the LCVR.

By measuring the intensities at 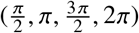 retardances for the PSA-LCVR, we obtain a system of 4 equations with 3 Stokes vector components, which are solved for *s*_0_, *s*_1_, and *s*_3_. The obtained values are averaged to find a unique value for each of the components [42]. Note, that *s*_2_ of SHG signal component can be measured with the addition of a second LCVR on the PSA. The SHG Stokes vector components *s*_0_, *s*_1_, and *s*_3_ are obtained for 4 incident polarization states having PSG retardances at *θ* = *π*, 2*π* for orthogonal linear polarizations VLP and HLP, and analogously, 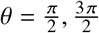 for the orthogonal circular polarizations LCP and RCP, respectively. The incident polarization states (e.g HLP, VLP, LCP, RCP) will be indicated in the superscript for each Stokes vector component.

Several polarimetric parameters are extracted to characterize the tissue. The polarimetric parameter can be expressed in terms of achiral *R* and chiral *C* susceptibility component ratios for the *C*_6_ symmetry model, and are defined as follows [46]:

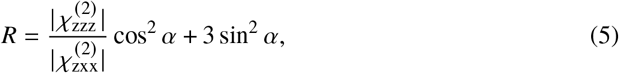

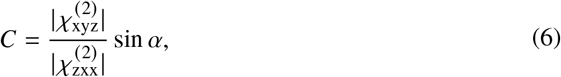

where 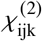 represents the ijk molecular component of the second order susceptibility tensor and α corresponds to the out-of-image-plane tilt angle of the collagen fibrils.

For in-plane orientation-independent assessment of the tissue ultrastructure the circular incident polarizations can be employed. Consequently, the SHG circular dichroism is expressed as [15, 38, 47]:

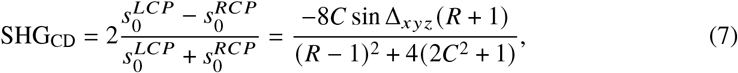

where Δ_*xyz*_ corresponds to the relative phase of the chiral 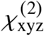 component with respect to the achiral components [48]. Note alternate definition of polarimetric parameters in [48] and in this work. It can be seen that SHG_CD_ depends on the C-ratio that can flip the sign depending on the alpha orientation angle (see Eq. 6). The SHG_CD_ also depends on the phase shift Δ_*xyz*_ between achiral and chiral components, while achiral susceptibility components are considered to be in phase [48]. The SHG_CD_ also depends on the R-ratio, but the sign of SHG_CD_ will not flip due to *R* having the quadratic dependence on sin *α* and cos *α* (see Eq. 5). Circular anisotropy of circular dichroism of SHG (CA_CD_) can be measured using circular polarization states of the PSA, expressed as:

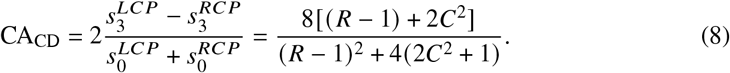

The out of plane angle U dependency is explicitly included in the R- and C-ratios. Moreover, C-ratio has a quadratic dependence for CA_CD_, and when assumed to be small, it can be dropped out of the equation. Then the R-ratio can be calculated [15]:

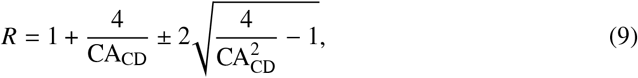

where the physically adequate range of *R* defines the correct sign. In addition, the degree of circular polarization (DCP) can be defined as follows [15, 49]:

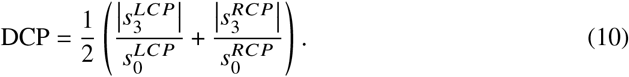

We can also obtain the in-plane orientation-independent average SHG intensity when the incident circularly polarized states are employed. The intensity is expressed in terms of Stokes vector components, namely:

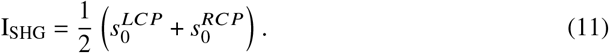

Notice that I_SHG_, SHG_CD_, CA_CD_, and DCP parameters are independent of the in-plane angle of the fibers, *δ*. If in plane orientation *δ* of the fibers is of interest, polarimetric parameters involving linear polarizations can be employed. The SHG linear dichroism (SHG_LD_) can be expressed as follows:

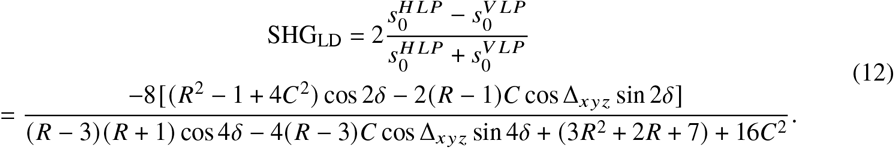

It can be noted that the numerator of SHG_LD_ varies with 2*δ* modulation frequency, while denominator is dependent on 4*δ* modulation. In addition, the C-ratio introduces the phase shift for cos 2*δ* modulation. Therefore, SHG_LD_ image will exhibit only approximate in-plane orientation of the fibers with bigger deviation for pixels with larger *C* cos Δ_*xyz*_ values. The sign of *C* cos Δ_*xyz*_ will also play a role in the deviation direction from cos 2*δ* dependence.

### 2.4. Sample preparations

The bleaching kinetics were measured from H&E stained and unstained Achilles tendon, harvested from a 25 kg Yorkshire pig euthanized within another approved research study at University Health Network, Toronto, Canada. The tendon was fixed in 10% buffered formalin and cut longitudinally along the tendon axis. The samples were embedded in parafin, and sectioned into 5- *μm* thick slices. The slices were mounted on microscope slides and used unstained, or were stained with hematoxylin and eosin.

A tissue microarray slide was assembled containing human lung, breast, kidney and liver tissue cores. The tissues were obtained according to an institutionally-approved protocol (University Health Network, Toronto, Canada). The tissue cores were 0.6 mm in diameter composed of formalin-fixed tissue specimens sectioned into 5 *μm* thick slices and mounted on a glass slide. The sections were stained with hematoxylin and eosin and imaged with a bright-field microscope scanner (Aperio Whole Slide Scanner, Leica Biosystems) for reference. Assessment of the tissue architecture was performed by an expert pathologist (S.J.D.).

## 3. Results and Discussions

### 3.1. Wide-field microscopy imaging conditions for H&E stained histology sections

The wide-field multiphoton excitation microscopy requires assessment of imaging conditions for biological samples, in particular, with respect to photobleaching of fluorescence and SHG signals. The effect of laser beam intensity and pulse repetition rate on the bleaching of fluorescence and SHG signal was investigated by imaging H&E-stained pig Achilles tendon tissue. Typical fluorescence and SHG images of a pig tendon before and after 10 minutes of wide-field illumination are presented in Fig 2a,b and c, d, respectively. The bleaching of fluorescence is noticeable, while SHG intensity only slightly increases after illumination. It demonstrates that imaging with SHG can be conducted for a long period of time, but MPF has to be checked for photobleaching.

**Fig. 2.**
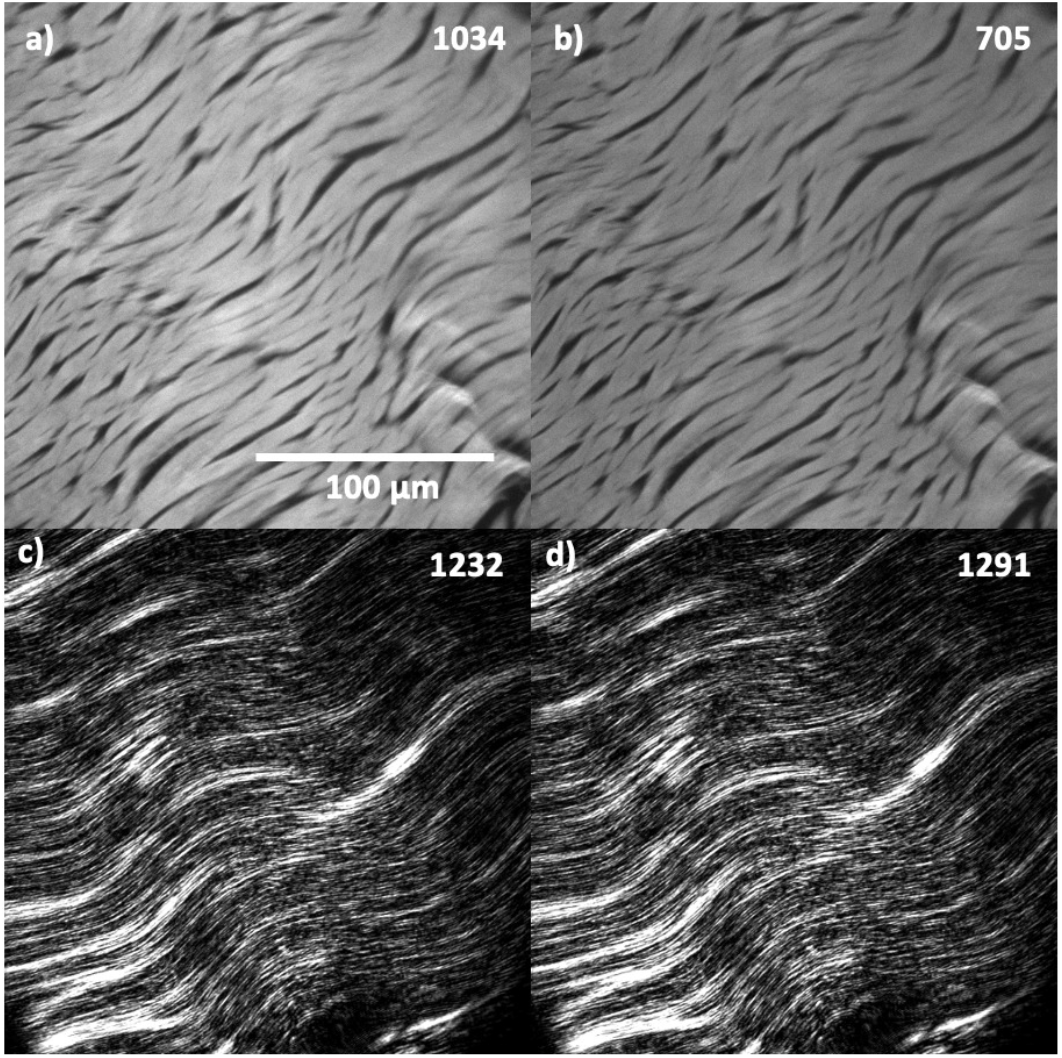
Signal bleaching for MPF (a,b) and SHG (c,b) images with a 10 minute interval between the images. A pulse energy per area of 0.014 J/cm2 was used for MPF. SHG signal in the same time interval is shown in panels (c) and (d), using a pulse energy per area of 0.017 J/cm^2^. Maximum pixel intensity is shown in the upper right side of each image. Notice, SHG signal doesn’t bleach, instead it increases slightly. All the measurements were done with a laser repetition rate of 1MHz

Detailed investigations were carried out to assess the characteristics of photobleaching in H&E stained histology tissue slices. The tissue was continuously illuminated for 20 minutes and images were taken every minute during the exposure. The time dependent bleaching curves for fluorescence and SHG signals were constructed for different laser pulse energies per area and different pulse repetition rates, as shown in Fig. 3. The average pixel value for each acquired image was chosen for the bleaching analysis. Selected time-dependent fluorescence and SHG signal curves are shown in Fig. 3a. It can be seen that fluorescence was bleached by 88% during the 20 minutes exposure with high pulse energy per area of 0.046 J/cm^2^ at 275KHz repetition rate, while SHG signal increased during that time period, using the same laser parameters. The kinetics had different rates for the fluorescence and SHG signals. The time-dependent increase in SHG in the stained samples could be in part attributed to the decrease in reabsorption of SHG signal due to bleaching of eosin. The time-dependent autofluorescence and SHG signals were also measured for the unstained sample (Fig. 3a). In the unstained sample, SHG intensity remains approximately constant over time. In contrast, the autofluorescence bleaching is evident. The autofluorescence originates mainly from flavin mononucleotide (FMN) and flavin adenine dinucleotide (FAD) molecules at 1030nm excitation [50, 51]. Therefore, when H&E-stained histology section is imaged, there is more than one fluorophore involved in the fluorescence, and different fluorophores bleach with different rates leading to the multiexponential bleaching kinetics.

**Fig. 3.**
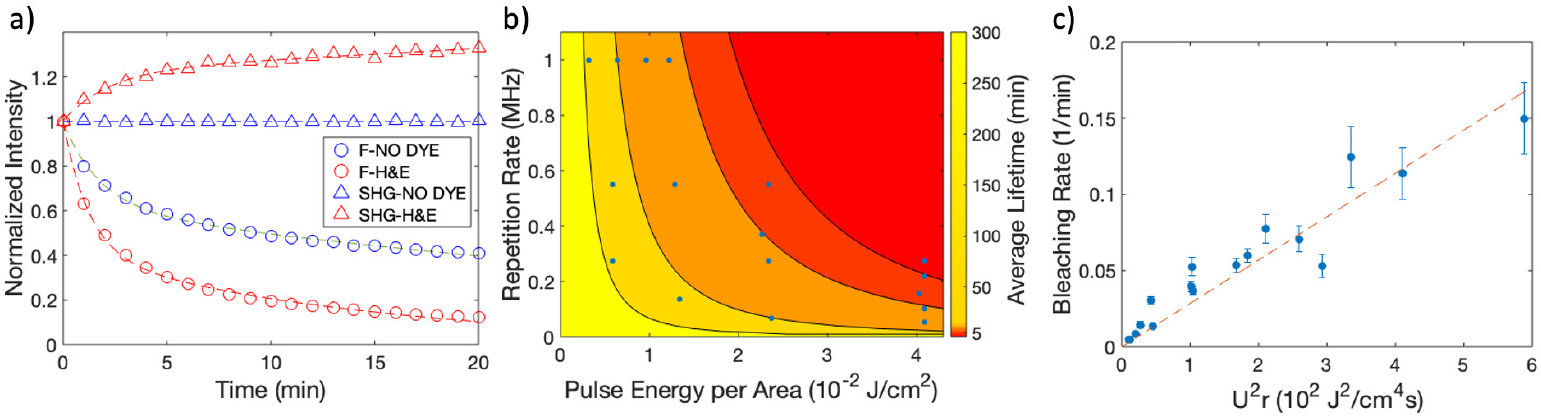
Fluorescence and SHG signal bleaching analysis for wide-field illumination with different pulse energies per area and at various pulse repetition rates. a) Normalized intensity of fluorescence (circle) and SHG (triangle) signal change over time for H&E stained (red) and unstained (blue) samples. Stained samples where imaged with a pulse energy per area of 0.046 J/cm^2^ at 275KHz repetition rate. Unstained samples were imaged with pulse energy per area of 0.025 J/cm^2^ at 550KHz repetition rate. b) Contour plot of average fluorescence bleaching lifetime dependence on pulse energy per area and pulse repetition rate. The data (blue dots) were globally fit with Eq. (3). c) The MPF bleaching rate dependence on pulse energy per area squared times pulse repetition rate.

The bleaching rate dependence on the laser parameters was assessed by performing a double exponential fit to the MPF bleaching decays with Eq. (2). The calculated average bleaching rate values at different pulse energies per area and pulse repetition rates were fit with Eq. (3), while keeping the exponent of *U*^*m*^ as a fitting parameter. Assuming a linear dependence of bleaching rate on the pulse repetition rate, the fit of the bleaching rate dependence on the pulse energy per area gave an exponent of *m* = 2.041 ± 0.059, which is in agreement with the two-photon excitation process for the fluorescence. Hence, the result is consistent with Eq. (3), and, therefore, validates the two-photon excitation fluorescence bleaching model presented in the Appendix. The fitted data in the form of a contour plot of fluorescence bleaching lifetime (1 / Γ) dependence on the pulse energy per area and pulse repetition rate is presented in Fig. 3b. The blue dots represent the average bleaching lifetimes obtained from the measured fluorescence bleaching kinetics at corresponding pulse energies per area and pulse repetition rates. The surface plot can be used to estimate the fluorescence bleaching lifetime for selected pulse energy per area and pulse repetition rate. It is evident that experiments with minimal bleaching can be performed for the laser beam parameters falling into the yellow/reddish zone of the contour plot in Fig. 3b.

To check if two-photon excitation model works well for all chosen pulse energies per area and pulse repetition rates, a linear fit of *U*^2^r dependence on the bleaching rate is presented in Fig. 3c. The linear fit to the data (see Eq. 3) shows that all experimental values follow the linear dependence, indicating that model works even at high pulse energies per area or high (1 MHz) pulse repetition rates. There are no indications of additional bleaching processes e.g due to heating or due to higher order photon absorption processes. Therefore, the photobleaching effect can be well estimated over a broad range of laser parameters using two-photon excitation model. It can be seen from Fig. 3b that fluorescence bleaching can be kept to a minimum, for example, with average bleaching lifetime of 20 min (yellow/reddish zone), for laser pulse energies per area below 0.03J /cm^2^ and a pulse repetition rate of 125 KHz, i.e. the average power density below 3.75KW /cm^2^. For pulsed lasers with a higher repetition rate, namely 1 MHz, imaging with the same bleaching lifetime (20 min) can be achieved below 0.01J /cm^2^ energy per pulse per unit area i.e. the average power density below 10KW /cm^2^. Such laser setings can be used for experiments with sample illumination time lasting for several minutes. If longer illumination times are needed, the laser power density has to be reduced. Note, that SHG signal is much less affected by the bleaching process and leads to a slight signal increase during the measurements as mentioned above (Fig 2).

It is also interesting to note that according to Eq. (1), the same dependence on pulse energy per area and the repetition rate is in the pre-exponential factor and also in the bleaching rate expression. Therefore, both the initial fluorescence intensity and bleaching rate will be affected by the same factor depending on the laser beam parameters. In practice, the fluorescence bleaching rate is rather slow, with characteristic lifetime on the order of tenths of minutes, which allows for prolonged imaging of the samples and can be used for polarimetry with a number of different polarization states.

### 3.2. Wide-field polarimetric SHG microscopy in histopathology samples

An application of wide-field polarimetric SHG microscopy for histology studies is presented in Fig. 4. The wide-field polarimetric microscopy was applied for imaging lung, liver and kidney tissue cores from the tissue microarray. The imaging was performed with minimized fluorescence and SHG bleaching conditions. Entire cores of the tissue, 600 ’m in diameter, were polarimetrically imaged using a sCMOS camera with 10 seconds exposure time. The 16 combinations of polarization states were recorded. The presented kidney tissue possessed the largest collagen content as seen in the SHG intensity images in Fig. 4. The SHG images were acquired with circularly polarized light, see Eq. 11. The R-ratio varied slightly between the tissues. The average R-ratio values for kidney, liver and lung were 1.90 ± 0.33, 2.02 ± 0.29 and 1.85 ± 0.35, respectively. The R-ratio reflects the ultrastructural organization of the collagen and can be used in cancer diagnostics [12, 14, 16, 17]. The peak of DCP value distribution was somewhat similar for all three tissues, however, liver DCP distribution is narrower, partly due to low amount of collagen in the tissue. The distribution of SHG_CD_ to positive and negative values shows that collagen orientation with out of image plane tilt in the tissues is widely distributed (see Eqs. 6, 7). The SHG_CD_ is also influenced by the distribution of molecular achiral and chiral susceptibility ratios (Eqs. 5, 6 and 7) and the relative phase Δ_*xyz*_. The wide distribution is also present in the SHG_LD_ map, indicating broad in-image-plane orientation distribution of angle *δ* values of collagen fibers (Eq. 12). The SHG_LD_ distribution is also influenced by the R-and C-ratios and the relative phase Δ_*xyz*_. The polarimetric parameters can be used for ultrastructural characterization of the histology tissue and for cancer diagnostics [42].

**Fig. 4.**
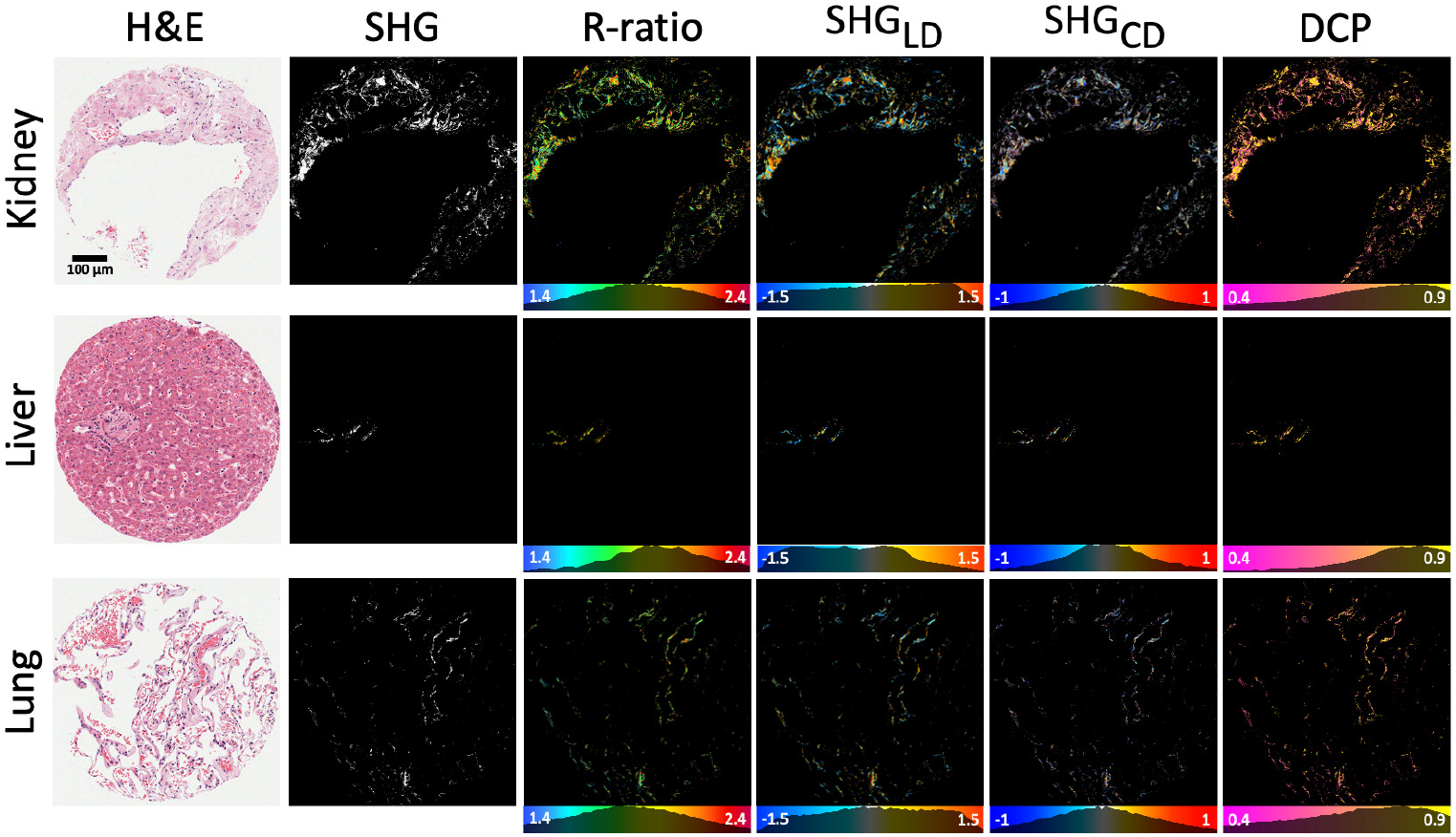
Images of polarimetric SHG parameters for kidney, liver and lung tissue cores from the tissue microarray. The respective rows represent images for kidney, liver and lung. The first column represents white light images of H&E stained core sections. The second column represents SHG intensity images. The other respective columns represent maps of calculated polarimetric parameters of R-ratio, SHG_LD_, SHGCD, and DCP. The color bars below the images contain the occurrence histograms of pixel values.

**Fig. 5.**
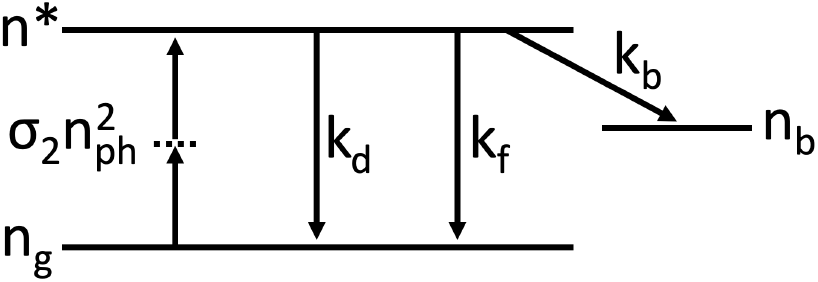
The dye excitation and relaxation processes occurring during illumination. The *n*_*g*_, = *n\** and *n*_*b*_ denote number of molecules in the ground state, excited state and bleached state, respectively. The σ_2_ denotes the two-photon exitation rate, and *n*_*ph*_ is the number of excitation photons. The *k*_*f*_, *k*_*d*_ and *k*_*b*_ represent radiative decay, non-radiative decay and bleaching rates, respectively.

Wide-field SHG microscopy enables fast imaging of histology samples over large areas. An imaging time for one polarimetric state was 10 seconds and the whole polarimetric set of 16 states was imaged in 8 minutes. The following laser parameters were used for the imaging (Fig 4): repetition rate of 100 KHz and 0.31 mJ/cm^2^ pulse energy per area. During the 8 minute long measurement, the fluorescence signal bleached by < 5%, while the SHG signal increased by less than < 1%. Therefore, the bleaching effects are within the measurement errors for different polarization states in the experiment. The bleaching effect for SHG can be further reduced with imaging unstained tissue. With the wide-field polarimetric SHG microscopy, the imaging time of large area is markedly reduced compared to the laser scanning mode. Therefore, the application of wide-field nonlinear microscopy for histopathology enables the whole slide scanning in a reasonable time, with minimal photodamange and photobleaching effects. This brings multimodal MPF and SHG microscopy closer to the clinical applications of digital pathology.

## 4. Conclusions

Bleaching kinetics were theoretically modelled and fitted using different experimental parameters, such as pulse energy per area and pulse repetition rate. We obtained experimentally that the square of the pulse energy per area is proportional to the bleaching rate; this is in agreement with fluorescence bleaching being mainly a two photon excitation process. Optimal ranges for laser parameters have been determined in order to have efficient wide-field polarietric SHG imaging, resulting in minimal MPF bleaching, i.e. less than 5% after 8 minutes of illumination and less than 1% SHG signal increase for H&E stained samples. For a low repetition rate lasers (125KHz) it is recommended to use pulse energies per area below 0.03J /cm^2^. Alternatively, for pulsed lasers with a higher repetition rate (1MHz), energy per pulse per unit area should be kept under 0.01J /cm^2^.

Wide-field nonlinear microscopy has been successfully employed for histopathology tissue imaging. The demonstrated SHG imaging of 700’< x 700’< sample area was accomplished with 1s exposure time. Wide-field nonlinear microscopy enables whole-slide imaging in less than 3 minutes, bringing a new era for nonlinear digital pathology investigations.

## Appendix

### 4.1. Nonlinear excitation fluorescence bleaching kinetics

The bleaching kinetics of multiphoton excitation fluorescence are modeled by assuming photo-induced accumulation of non-fluorescing chromophores without further quenching of the fluorescing molecules. The fluorescence dye (eosin) contained in the tissue is excited by a two-photon excitation process, and undergoes relaxation that includes radiative and non-radiative decay to the ground state. In addition, the excited singlet state can decay to a bleached state. Usually this occurs when excited molecule undergoes intersystem crossing into a triplet state and further into a dark state, for example, via radical formation or reaction with oxygen. Therefore, by converting to the dark state chromophores get out of the pool of fluorescent molecules. In this model, the bleached dye molecules reduce the concentration of fluorophores in the ground state and do not participate in fluorescence quenching. The scheme of excitation and relaxation processes is presented in Fig.5.

The rate for generation of excited states of the dye molecules is:

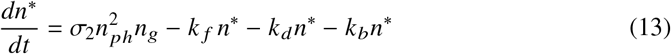

where *n\** is the number of molecules in excited state, *n*_*g*_ the number of molecules in ground state, and *n*_*ph*_ is the number of photons available for two-photon excitation of molecules. The two-photon excitation rate is σ_2_, and *k*_*f*_, *k*_*d*_ and *k*_*b*_ are the rate constants for radiative decay, nonradiative internal conversion and bleaching, respectively. The time is represented by t.

The time-dependent change in the number of dye molecules in the ground state is represented by the following equation:

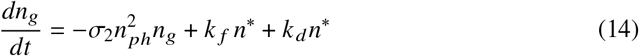

It can be seen that depopulation of the ground state occurs due to nonlinear absorption, and repopulation due to the excited state decay to the ground state via radiative and non-radiative decay. The generation of bleached molecules can be described by the reaction from excited state with the rate constant *k*_*b*_:

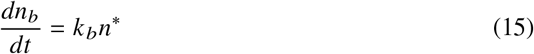

In addition, we will use a balance equation for the dye molecules, where total number of molecules is equal to sum of molecules in the ground state, excited state and bleached state:

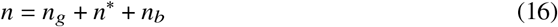

The excitation process and the relaxation processes to ground state are much faster than the transition to bleached state with the rate constant *k*_*b*_. Therefore, 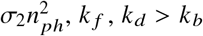, and thus we consider that molecules in the excited state reach a quasi-steady-state equilibrium, rendering *dn*\*/ *dt* = 0, and:

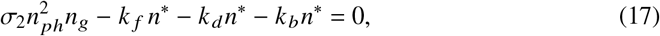

which gives

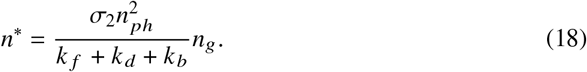

The number of fluorophores in the ground state is:

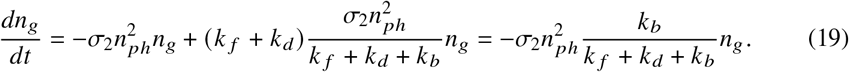

After integration the above equation gives:

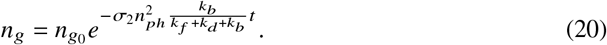

It can be seen that number of molecules in the ground state exponentially decreases from the initial value *n*_*g*0_. The fluorescence intensity *I*_*f*_ is proportional to the number of excited molecules and the radiative rate:

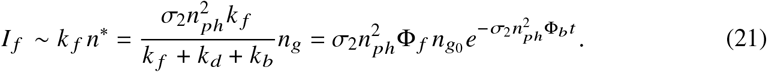

where the fluorescence yield is expressed 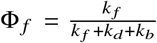 and the bleaching yield is 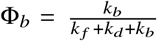. It can be seen that fluorescence intensity will decrease over time with the exponential decay. The fluorescence bleaching rate depends on the number of photon pairs, for two-photon excitation process. The number of photon pairs will depend on the squared pulse energy per unit area, *U*^2^, and the repetition rate of the laser, 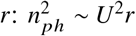. By including this dependency into the fluorescence decay equation, we have:

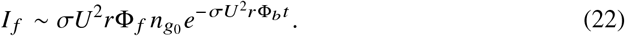

The fluorescence bleaching kinetics can be analyzed by normalizing fluorescence decay equation to the initial fluorescence intensity, *I*_0_:

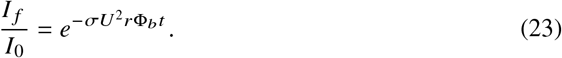

The fluorescence bleaching curve can be examined for the dependence on laser pulse energy per unit area and pulse repetition rate. The fitted fluorescence bleaching kinetics can determine the σ Φ_*b*_ parameter. The product of two-photon absorption rate and bleaching yield will have a common value for decays at different pulse energies per unit area and pulse repetition rates. Drastic variations of extracted σ Φ_*b*_ value for different laser pulse energies and repetition rates, would indicate the presence of additional processes involved in the bleaching.

